# ARAX: a graph-based modular reasoning tool for translational biomedicine

**DOI:** 10.1101/2022.08.12.503810

**Authors:** Amy K. Glen, Chunyu Ma, Luis Mendoza, Finn Womack, E. C. Wood, Meghamala Sinha, Liliana Acevedo, Lindsey G. Kvarfordt, Ross C. Peene, Shaopeng Liu, Andrew S. Hoffman, Jared C. Roach, Eric W. Deutsch, Stephen A. Ramsey, David Koslicki

## Abstract

**Motivation:** With the rapidly growing volume of knowledge and data in biomedical databases, improved methods for knowledge-graph-based computational reasoning are needed in order to answer translational questions. Previous efforts to solve such challenging computational reasoning problems have contributed tools and approaches, but progress has been hindered by the lack of an expressive analysis workflow language for translational reasoning and by the lack of a reasoning engine—supporting that language—that federates semantically integrated knowledge-bases.

**Results:** We introduce ARAX, a new reasoning system for translational biomedicine that provides a web browser user interface and an application programming interface. ARAX enables users to encode translational biomedical questions and to integrate knowledge across sources to answer the user’s query and facilitate exploration of results. For ARAX, we developed new approaches to query planning, knowledge-gathering, reasoning, and result ranking and dynamically integrate knowledge providers for answering biomedical questions. To illustrate ARAX’s application and utility in specific disease contexts, we present several use-case examples.

**Availability and Implementation:** The source code and technical documentation for building the ARAX server-side software and its built-in knowledge database are freely available online (https://github.com/RTXteam/RTX). We provide a hosted ARAX service with a web browser interface at arax.rtx.ai and a web application programming interface (API) endpoint at arax.rtx.ai/api/arax/v1.3/ui/.

## 1 Introduction

Structured biomedical knowledge is rapidly accumulating in primary databases such as ChEMBL [1], DrugBank [2], Reactome [3], OBO Foundry [4], KEGG [5], Semantic MEDLINE Database (SemMedDB) [6], UniProtKB [7], and the Unified Medical Language System (UMLS) [8]. Using knowledge-based computational reasoning to answer translational questions such as *What drugs might be repurposed to treat Adams-Oliver Syndrome?* requires integrating facts spanning a variety of concept types including drugs, targets, pathways, genetic variants, phenotypes, and diseases. To facilitate such computational reasoning, there have been numerous efforts to integrate knowledge from various biomedical databases using knowledge graph (KG) abstractions [9, 10, 11] consisting of a labeled multigraph in which each node represents a concept and each edge represents a concept-concept relationship, i.e., a “triple”. Several such biomedical KGs have been described [12, 13, 14, 15, 16, 17, 18], including RTX-KG2, which we developed and described previously [19] and which integrates all of the aforementioned primary databases with a semantic layer that is described by the open-standard Biolink model [20]. Given such a comprehensive KG with a unified semantic layer, a biomedical question can be transformed into a *query graph* [21] (which represents a search pattern, analogous to a `select` statement in the SPARQL language [22]; see Fig. 1 for more details) and/or a *graph analysis workflow* in order to generate a list of “answers”/”results”, each corresponding to a *subgraph* of the KG. To facilitate that process for translational applications, an open-source, web-based, and standards-based tool is needed for designing, expressing, executing, and refining KG analysis workflows.

**Figure 1:**
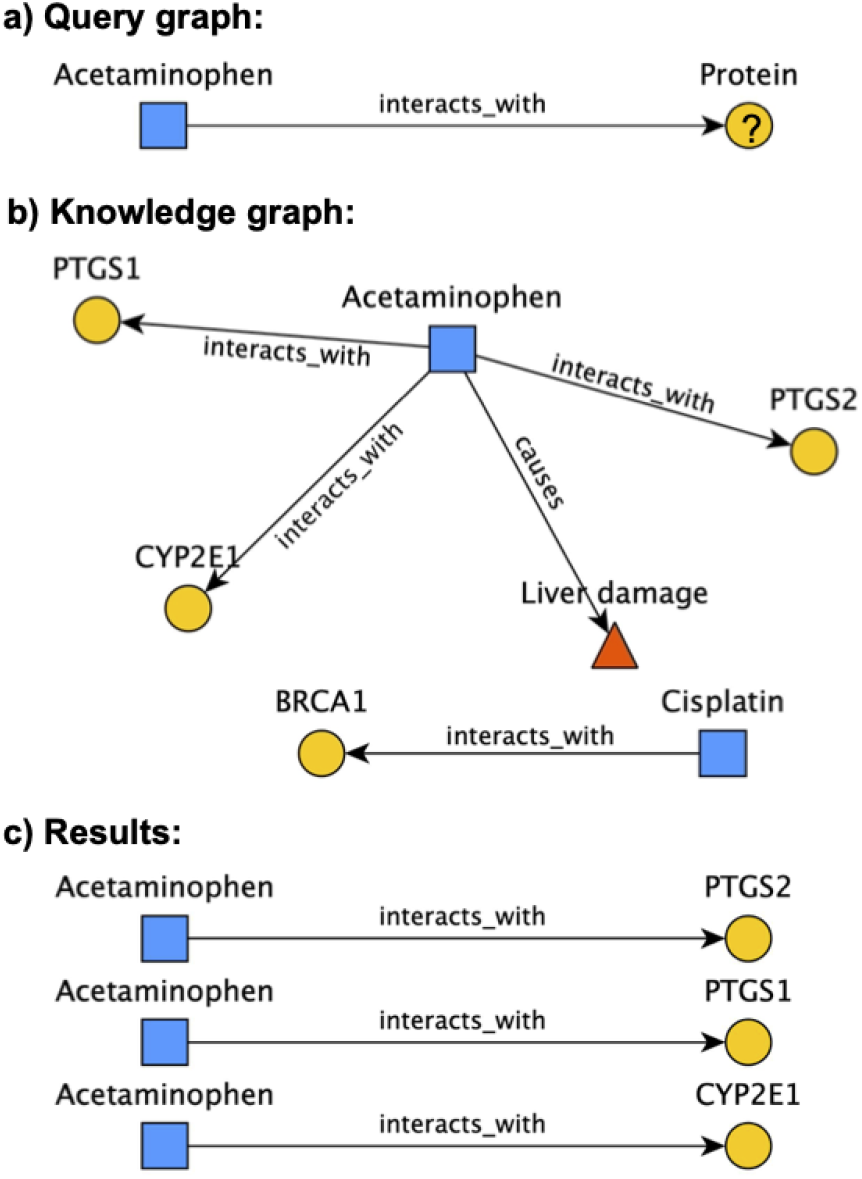
An example query graph (a) and the set of subgraphs that “fulfill” it (c) from an example knowledge graph (b). **Item a** depicts a query graph that asks for proteins that interact with the drug acetaminophen. Query graphs are connected graphs that define particular *patterns* to search for in graph data. In this example, the acetaminophen query node is considered “pinned”, because it is constrained to a specific concept; Biolink-defined [20] node categories (`Protein`) and edge predicates (`interacts_with`) constrain the rest of the query. While this query graph is quite simple, there is no hard limit to how many nodes/edges query graphs can include (though there may be a practical limit, due to the tendency for query runtime and memory consumption to increase with query graph size), and they may contain cycles and/or be multigraphs. **Item b** depicts a hypothetical knowledge graph (KG) that might be used to answer this query. This KG contains three different categories of nodes (represented by node shape/color) and two different edge predicates. **Item c** shows the three subgraphs of the KG that fulfill the query graph. A subgraph fulfills the query graph if it fits the pattern of the query graph in terms of its structure and constraints, with acetaminophen connected to a protein via an `interacts_with` edge. This example is simplified for clarity’s sake, but in reality there are more complexities to what it means to fulfill a query graph. For instance, Biolink node categories and edge predicates are hierarchical in nature, meaning a valid result for the example query graph could, for instance, use the edge predicate `physically_interacts_with`, because that predicate is a descendant of interacts_with. In addition, one could specify *multiple* node categories or edge predicates on a single node/edge in their query graph; in that case, nodes/edges with *either* of those categories/predicates are considered to fulfill that query node/edge.

Efforts to develop a comprehensive platform for expressing, executing and refining KG analysis workflows have used new query languages, web application programming interface frameworks, computational reasoning paradigms, graph topology heuristics, and machine-learning methods. BIOZON [23] provides a SQL-based query and search function for its KG; Hetionet [14] integrates multiple databases into a single graph for drug repurposing and leverages graph path-finding; BioGraph [24] uses the Gremlin graph query language; SPOKE [25] provides capabilities for finding a concept of interest and visualizing its “node neighborhood”; BioThings Explorer [26] builds a biomedical knowledge graph by querying SmartAPI-registered APIs; ROBOKOP [16] constructs a query-specific knowledge graph which it analyzes to identify candidate answers; and mediKarnren [27] uses the miniKanren logic programming system for drug repurposing applications. Despite much progress, two key problems remain: (1) how to provide control and transparency of the graph analysis workflow that is executed to answer a user’s question and (2) how to rank potential answers. The control/transparency problem entails a trade-off: on one extreme, a general-purpose graph query language such as Neo4j Cypher^1^ provides complete control, but expressing, modifying, and extending a state-of-the-art drug repositioning analysis workflow is extremely complex and demanding for many users. On the other extreme, canned analysis workflow approaches in practice have limited customizability. In addition, many tools have a limited number of knowledge sources and/or lack a standardized semantic layer with sufficient expressiveness.

### 1.1 ARAX overview and key advantages

Here we describe ARAX (arax.rtx.ai), a new computational reasoning tool for querying and exploring biomedical knowledge and data. ARAX provides three key advantages versus previous systems:

1. ARAXi, ARAX’s intuitive language for specifying a workflow for analyzing a knowledge graph (Sec. 2.1);
2. access to around 40 knowledge providers (which themselves access over 100 underlying knowledge sources) from a single reasoning tool, using a standardized interface and semantic layer (Sec. 2.2); and
3. a versatile method for scoring search result subgraphs (Sec. 2.1.5).

Together, these advantages are designed to facilitate and improve reasoning-based answering of translational questions [28] such as *What genetic conditions protect against asthma? What drugs target proteins associated with the cyclooxygenase pathway?* and *How can expression of KCNMA1 be pharmacologically inhibited?*

ARAX is one of six reasoning engines in the Biomedical Data Translator system (henceforth, “Translator”) [29, 30], (see “Funding”), a distributed computational system for accelerating translational science. Translator has a layered architecture; queries are interpreted by a coordinating service, which sends the query to the reasoning engines which, in turn, consult registered Knowledge Provider (KP) services to answer the query. All Translator services exchange messages using a standard web interface, the Translator Reasoner API (TRAPI),^2^ whose technical specification is collaboratively maintained. For its semantic layer, TRAPI uses the Biolink Model^3^ to specify types of biological entities and relationships. ARAX can be directly queried via either its TRAPI API or its web browser user interface; these provide various ways to formulate queries and explore results (Sec. 2.3).

A typical query and corresponding workflow ARAX may use to answer it is depicted in Figure 2, along with a diagram of ARAX’s architecture. Its architecture is modular, with five different analysis modules that can be run using ARAXi commands (Sec. 2.1). ARAX is mainly built on top of RTX-KG2 [19], our large-scale biomedical knowledge graph that has a TRAPI-compliant query interface and is registered as a Translator KP; but ARAX also queries close to 40 other KPs with specific areas of emphasis (Sec. 2.2.1), including genome-wide association study data, electronic health record data from medical centers, and molecular data from high-throughput cellular assays.

**Figure 2:**
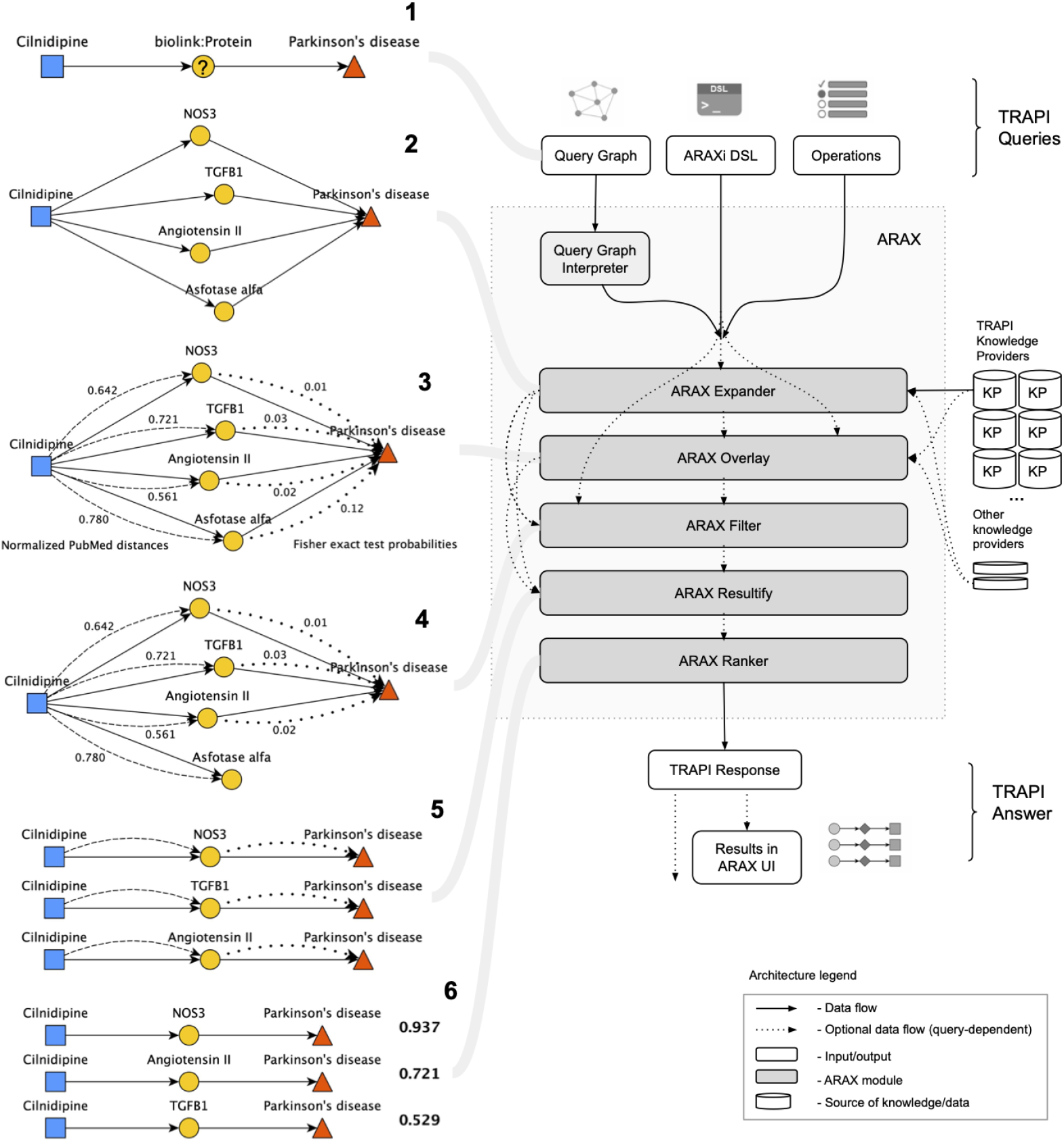
ARAX answers queries using *workflows*, in which each step is handled by a particular ARAX module. **Left panel:** The left half of this figure depicts a typical simple query that might be submitted to ARAX (Step 1) and how ARAX goes about answering that query (Steps 2-6). More specifically, Step 1 shows a visual representation of the query in query graph form, which asks for proteins associated with both Parkinson’s disease and the drug cilnidipine. Step 2 depicts the Expand step, which reaches out to various knowledge providers for answers to the given query and combines their answers into an answer knowledge graph. Step 3 represents the Overlay step, in which the answer KG is overlaid/annotated with contextual quantitative information in “virtual” edges (dashed/dotted lines). Step 4 depicts the Filter step, in which nodes and edges are filtered from the answer KG based on the statistical information overlaid in Step 3. Step 5 represents the Resultify step, which finds all of the subgraphs in the answer KG that fulfill the query graph. Step 6 represents the Rank step, which calculates a score for each result subgraph indicating the overall degree of epistemic support for that answer. **Right panel:** The right half of this figure depicts ARAX’s architecture, which is centered around five modules, each of which corresponds to a typical workflow step. Notably, ARAX accepts queries in three different forms, all of which are based on the Translator Reasoner API (TRAPI) data model: 1) a query graph (as shown in Step 1), 2) the ARAXi graph analysis workflow definition language (Sec. 2.1), or 3) TRAPI Operations. The ARAX system translates the two non-ARAXi input forms into an ARAXi workflow that specifies the series of ARAX modules that will be run to answer the query. The five core ARAX modules (Expander, Overlay, Filter, Resultify, and Ranker) operate independently and may be combined in various orders depending on the query. When the input is in the form of a query graph, the Query Graph Interpreter layer selects an appropriate series of ARAXi commands to answer the question at hand; this selection of ARAXi commands is done using a template-based system in which templates were manually curated by examining common query graph structures or archetypes. The final answer knowledge graph and results are returned in a TRAPI JavaScript Object Notation (JSON) response either directly to the requesting software program or to the ARAX UI for browser rendering (Sec. 2.3.1).

As illustrated in Figure 2, ARAX accepts input queries in any of three representations: a TRAPI query graph, ARAXi, or TRAPI Operations. TRAPI query graphs are graph-based templates representing the user’s question (see Fig. 1), expressed in JSON format. The ARAXi language, which can be used to express a custom graph analysis workflow including overlaying of annotations and/or filtering of edges to reduce the result-set size, is described in Section 2.1. TRAPI Operations^4^, the third way of specifying a graph analysis workflow, is an ARAXi-inspired, Translator-standardized vocabulary; it is translated into ARAXi by the ARAX interpreter.

## 2 Methods

### 2.1 ARAXi knowledge graph analysis language

To address the need for an intuitive language for expressing KG analysis workflows, we developed ARAXi, a procedural language that allows a user or software tool to concisely express biomedical questions. The five main ARAXi modules (see Secs. 2.1.1–2.1.5) can be used individually or combined in a graph analysis workflow (Fig. 2). Each ARAXi command corresponds to an analysis module; this modular design simplifies both construction and reuse of knowledge graph analysis workflows.

All input queries—whether in the form of a TRAPI query graph, ARAXi, or TRAPI Operations—are ultimately translated into a series of ARAXi commands, which together define the knowledge-retrieval and analysis workflow for answering the query. In particular, two core ARAXi commands, `add_qnode` and `add_qedge`, are used to incrementally construct a query graph, which can then be expanded by the ARAX_expander module (Sec. 2.1.1) into a query-specific knowledge graph (KG); the query-specific KG can then be refined or have knowledge overlaid using the four other ARAX modules (Secs. 2.1.2–2.1.5).

#### 2.1.1 Expand a knowledge graph: ARAX_expander

The ARAX_expander module uses all TRAPI-compliant knowledge providers (KPs) registered in the SmartAPI registry [31] (see Sec. 2.2 for more details) to find subgraphs that satisfy the input query graph. For each edge in the query graph, ARAX_expander determines which KPs are capable of fulfilling that query edge (based on metadata provided dynamically by the KPs’ APIs) and then queries those KPs concurrently via their TRAPI-compliant APIs. When querying each KP, ARAX_expander converts any node identifiers in the query to semantically equivalent identifiers from controlled vocabularies that the KP prefers. This is accomplished by using the ARAX Node Synonymizer service (Section 2.2.2). ARAX combines answers returned from the KPs into a single answer knowledge graph that is “canonicalized,” meaning that it does not contain semantically redundant nodes.

ARAX_expander processes the query graph in a breadth-first fashion; that is, it retrieves all answers from KPs for a given query edge before moving on to the next query edge. It avoids combinatorial explosion by pruning down the answer set that it receives for each query edge in an adaptive and predictive manner: it uses the ARAX_overlay module’s Fisher Exact Test function (Sec. 2.1.2) combined with ARAX_ranker (Sec. 2.1.5) to decide which triples to retain for the given query edge. By default, the number of triples to retain for each query edge (the “pruning threshold”) is dynamically determined according to the projected degree of combinatorial explosion the given query edge will produce, based on heuristics pertaining to the depth of categories and predicates in the Biolink hierarchy. Currently these heuristics essentially set the threshold to one of three values when one of the query edge’s nodes is “unpinned” (not constrained to specific concepts): 100 if the given query edge is unconstrained in terms of categories and predicates or if the root Biolink categories/predicates are used, 200 if a category known to be very prevalent in KPs is used (this generally only includes categories one generation away from the root node in the Biolink category tree; e.g., `biolink:ChemicalEntity`), and 500 otherwise. Doubly-pinned queries are given a higher prune threshold of 5,000, due to their inherent constraint to specific concepts. Alternatively, a user may specify a per-query prune threshold that overrides the dynamically-determined default pruning via the ARAXi `prune_threshold` parameter. Having to prune answers after expanding each query edge is the main downside to the breadth-first approach of processing the query graph, since good answers may be lost; while a depth-first approach would avoid this, it allows for less batch querying of KPs (i.e., sending queries in which one node is pinned to many concepts) and thus results in prohibitively slow query times in our experience. In the future we plan to improve the dynamic pruning method to do query backtracking in some form; that is, if no answers for the next query edge are found based on the set of answers that were retained for the prior query edge, try using more of the prior query edge’s answers to see if they produce answers for the next query edge, and so on. This will effectively result in a sort of hybrid breadth-first/depth-first approach that may combine the best of both worlds.

Because users have different preferences in terms of the amount of time they are willing to wait for ARAX to provide answers to their query, ARAX_expander also provides a KP timeout parameter, which specifies how long ARAX_expander should wait for a response from each KP. Users who are more interested in getting answers quickly can specify a short (e.g., 30 seconds) timeout, while a user whose use-case prioritizes comprehensiveness over expediency can use a longer KP timeout.

#### 2.1.2 Overlay/annotate contextual information: ARAX_overlay

The ARAX_overlay module enhances the ARAX_expander-built KG by overlaying “virtual edges” (which are virtual in the sense that they do not fulfill edges present in the original query graph) to denote (1) structural similarity (i.e., a statistically significant number of shared neighbors) of two nodes in the KG; (2) a predicted “treats” relation between drug and disease nodes via a link-prediction model; or (3) statistically significant co-association in a database such as a clinical database, a combined clinical-epidemiological database, or the biomedical literature. Currently, the overlay function (which is called with the overlay command in ARAXi) can `overlay` nodes or edges with the following seven kinds of contextual information:

##### Fisher Exact Test

evaluates how significant/non-random a one-hop connection is between a given list of subject nodes and an object node based on the p-value calculated by following the traditional Fisher’s exact statistical test. This is equivalent to a phenotype or gene enrichment analysis in statistical genetics, but for arbitrary node categories. Given a set {*v_s_*} of subject nodes and an object node *v_o_*, a p-value is calculated to indicate how significant/non-random an edge is between the subjects and object. This requires using the following 2×2 contingency table as input to the “stats.fisher_exact” method in the Scipy python package [32]:

| | # nodes in $\{v_s\}$ | # nodes not in $\{v_s\}$ |
| --- | --- | --- |
| # nodes connected to $v_o$ | $a$ | $b$ |
| # nodes not connected to $v_o$ | $c$ | $d$ |

where “connected”/“not connected” means with respect to the knowledge graph RTX-KG2 when considering nodes of the same category as $v_o$ .
$\{a, b, c, d\}$ are the integer values representing number of nodes.

##### Jaccard Similarity

measures how many shared intermediate nodes with a specific category (e.g., “Protein”) can be found via knowledge sources between a subject node and an object node. For example, this can be used to find drugs that interact with many genes associated to a disease; find genes associated with many phenotypes of a disease; or other such “many intermediate connections” queries. More rigorously, given a starting node *v_s_* connected to *n* intermediate nodes 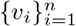 which each connects to an ending node *v_e_*, the Jaccard similarity between *v_s_* and *v_e_* is defined as follows, with OutDeg(v) representing the out-degree of a node in the knowledge graph:

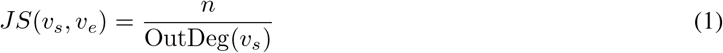

##### Drug-Treats-Disease Probability

predicts the probability that a given drug can treat a given disease. We developed this computational drug repurposing approach based on the model framework proposed by [33]. In order to adapt this model to our core knowledge graph RTX-KG2, we modified the implementation of the model by replacing node2vec [34] with GraphSage [35] for generating the node embeddings as well as using direct concatenation instead of the Hadamard product to generate the input features for a Random Forest classifier for drug-disease pairs. Additional performance gains may be realized through the use of alternative embedding or prediction models, though Section 3.3 indicates the current approach performs favorably.

##### Normalized PubMed Distance

measures the significance of co-occurrence of pairs of terms (corresponding to node names) in article abstracts in the PubMed (i.e., MEDLINE) database, using the “Normalized Google Distance” (NGD) measure [36]

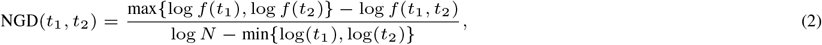

where *t*_1_ and *t*_2_ are the biomedical terms (node names) used in RTX-KG2; *f*(*t*_1_) and *f*(*t*_2_) respectively represent the total number of unique PubMed IDs associated with this term; *f*(*t*_1_, *t*_2_) is the number of unique and shared PubMed IDs between *t*_1_ and *t*_2_; and *N* is the number of pairs of article abstract and MeSH term [37] annotations in PubMed.

##### PubMed abstracts

obtained by mapping node identifiers to MeSH terms and obtaining the PubMed articles associated with this MeSH term.

##### Columbia Open Health Data (COHD) Clinical Information

[38] – obtained from electronic health records of 1.7M patients during a five-year period. For a given pair of terms, three statistical measures are obtained from COHD: observed clinical frequencies; the log-ratio between observed and expected counts; and the χ^2^ test statistic.

##### Exposure Data

statistical association data for pairs of terms in health records and between clinical terms and environmental exposure variables in epidemiological databases, obtained through the Integrated Clinical and Environmental Exposures Services (ICEES) service [39], which obtains data from University of North Carolina Health; the Drug Induced Liver Injury Network; and the National Institute of Environmental Health Sciences Personalized Environment and Genes Study.

#### 2.1.3 Filter the knowledge graph: ARAX_filter

Multi-hop queries can produce such large answer KGs that it is sometimes useful to prune down the KG to remove less important nodes or edges before proceeding. The ARAX_filter module allows one to do this; it provides several options for selectively removing nodes/edges based on their contextual information (often quantitative information added by ARAX_overlay) according to user-defined thresholds. This functionality can be invoked using the `filter_kg` ARAXi command. ARAX_filter also provides various methods for limiting the number of results returned and for sorting the results created by ARAX_resultify (see Sec. 2.1.4), for example, by score, edge/node counts, or particular edge/node attributes.

#### 2.1.4 Create results: ARAX_resultify

The ARAX_resultify module finds and returns all subgraphs in the answer KG that fulfill the query graph; this step is typically performed after the answer KG has been constructed and (optionally) pruned. ARAXi exposes a boolean `ignore_edge_direction` parameter that controls whether ARAX_resultify should ignore or enforce edge direction when determining whether a subgraph fulfills the query graph. When enforcing edge direction, ARAX_resultify will only return subgraphs for which edge directions match those of the query graph edges they are fulfilling. When ignoring edge direction, KG edges are allowed to fulfill query graph edges in the reverse direction (meaning, `PTGS2 - interacts_with -> Acetaminophen` would be a valid result for the query graph `Acetaminophen - interacts_with –> Protein`).

#### 2.1.5 Score and rank result graphs: ARAX_ranker

The ARAX_ranker module assigns a score between 0 and 1 (where a higher score represents a higher-confidence answer) to each result graph and orders the results by descending score. Scores are computed as follows:

1. For each “result graph”:
  a. Each edge in the result graph is assigned a score [0,1], as follows:
    i. Any scalar attribute (e.g., Fisher Exact Test p-value, or Jaccard Index) on the edge is normalized to a value between 0 and 1 using a sigmoid function parameterized for the specific type of attribute.
    ii. For any edges originating from SemMedDB, the attribute containing PubMed references for publications supporting the given assertion is converted into a scalar value via the function:

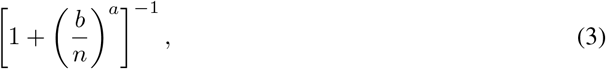

where *n* represents the number of publications listed in the attribute. The values of *a **=** log_2_**(9)*** and *b* = 4 were chosen so that 4 supporting publications results in a value of 0.5 (i.e. 50% confidence), and 8 publications gives a value of 0.9.
    iii. The given edge’s normalized, weighted scalar values are then multiplied to create one single score between 0 and 1 that is assigned to that edge.
  b. Using the normalized, weighted edge scores from Step 1a, three graph weights (calculated using max-flow, Frobenius norm, and weight-of-longest-geodesic-path) are obtained for the result graph.
2. Results are then ranked in descending order by their three metric scores calculated in Step 1b (in three separate lists).
3. Each result is assigned a final score that is the average of its three ranks from Step 2 (normalized between 0 and 1). The ARAX system automatically runs ARAX_ranker after ARAX_resultify, but ARAX_ranker can also be run individually via the rank_results ARAXi command; in principle, a list of result subgraphs produced by another Translator tool could be ranked using this ARAX service.

### 2.2 Knowledge/Data

#### 2.2.1 Richness

ARAX currently uses around 40 different Knowledge Providers to answer queries; the exact count depends on the number of TRAPI-compliant KPs registered in SmartAPI^5^ at query runtime, since ARAX_expander dynamically selects from all such KPs for a given query. The KPs themselves obtain information from various knowledge sources, giving ARAX access to a combined total of more than 100 knowledge sources (e.g., UniProt, DrugBank, DisGeNET, ChEMBL, MONDO, SemMedDB). Calculating the exact count of underlying knowledge sources is difficult due to the lack of such information in structured form for some KPs, but between RTX-KG2’s 73 knowledge sources [19], SPOKE’s approximately 40 knowledge sources^6^, and the BioThings Explorer Service Provider’s 32 integrated sources^7^, ARAX has access to at least 100 distinct underlying knowledge sources (we note that there is some overlap in knowledge sources between KPs). The true count of underlying knowledge sources is likely notably higher since ARAX uses many other KPs that were not included in the above estimate.

In total, ARAX’s KPs can answer queries about 135 different categories of nodes (e.g., `biolink:Protein, biolink:Disease`) and 281 different edge predicates (e.g., `biolink:interacts_with, biolink:treats`). This results in a total of 76,443 distinct meta-triples (combinations of subject node category, edge predicate, and object node category) that ARAX has access to. The breakdown of these counts by KP is shown in Table 1 for ARAX’s top 25 KPs in terms of number of supported meta-triples. Figure 3 shows a small selection of ARAX’s overall meta-graph, including some of the most commonly queried node categories. The counts for each KP are based on their TRAPI v1.3.0 API /`meta_knowledge_graph` endpoints, accessed on Dec. 12, 2022. All such APIs are listed in the SmartAPI Registry online 5.

**Figure 3:**
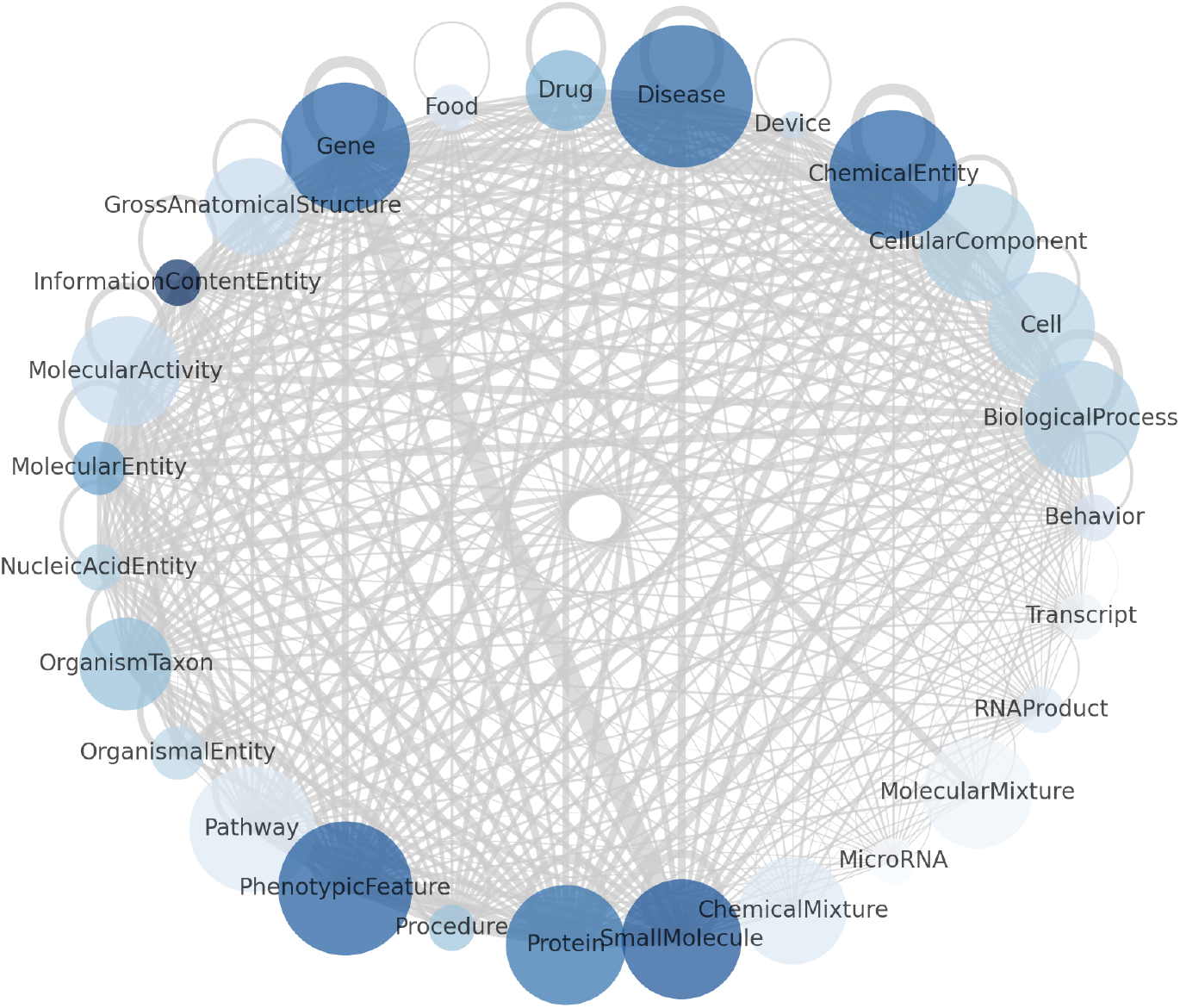
A small portion of ARAX’s meta-graph including a selection of the most commonly queried node categories. The size of each node represents the number of ARAX’s KPs that can answer queries involving that category, the color of each node represents the number of node identifier types ARAX has access to (through its KPs) for that category, and the thickness of each edge represents the number of distinct predicates ARAX has access to between two nodes with the given categories.

**Table 1:** ARAX’s top 25 knowledge providers in terms of number of meta-triples they can answer queries about. Abbreviations: Cat., Categories; Pred., Predicates; Meta., Meta-triples. More information on each KP along with links to their APIs are available in the SmartAPI registry (Footnote 5).

| Knowledge provider | Cat. | Pred. | Meta. |
| --- | --- | --- | --- |
| RTX-KG2 | 57 | 86 | 45,300 |
| CAM-KP | 105 | 50 | 28,601 |
| Ontology-KP | 81 | 14 | 4,434 |
| Automat-robokop | 20 | 190 | 2,296 |
| Automat-uberongraph | 18 | 66 | 1,414 |
| Service Provider (by BioThings Explorer) | 22 | 97 | 1,102 |
| Automat-ctd | 16 | 91 | 730 |
| SRI Reference KG | 14 | 89 | 644 |
| Text Mined Cooccurrence API | 18 | 1 | 322 |
| Automat-cord19 | 17 | 6 | 291 |
| MolePro (Molecular Provider) | 20 | 126 | 288 |
| Automat-biolink | 17 | 13 | 210 |
| Automat-hetio | 15 | 27 | 159 |
| Automat-ontology-hierarchy | 18 | 2 | 134 |
| Automat-drug-central | 11 | 17 | 104 |
| Automat-pharos | 13 | 12 | 95 |
| Automat-hmdb | 10 | 9 | 72 |
| Automat-human-goa | 10 | 23 | 60 |
| Automat-icees-kg | 8 | 1 | 57 |
| COHD (Columbia Open Health Data) | 5 | 2 | 50 |
| SPOKE KP | 18 | 24 | 44 |
| Automat-gtopdb | 6 | 11 | 40 |
| Automat-viral-proteome | 10 | 11 | 35 |
| Automat-panther | 8 | 7 | 24 |
| Automat-gtex | 2 | 20 | 20 |

#### 2.2.2 Knowledge Federation

Two aspects of ARAX’s architecture are crucial for knowledge federation: its adherence to standards for query APIs and for its semantic layer (i.e., TRAPI and Biolink, respectively) and its comprehensive service for mapping between equivalent node identifiers (i.e., node synonymization).

##### TRAPI and the Biolink Model

ARAX bypasses many of the data integration challenges that arise when combining information from multiple sources by adhering to a standard web API format (TRAPI ^2^) that itself adheres to a standard semantic format (Biolink^3^). Not only does ARAX’s API conform to TRAPI, but in the course of answering queries, ARAX largely only uses KPs that themselves speak TRAPI^8^. This means that ARAX has to do very minimal processing to relay queries to and combine answers from different KPs, since they all represent their answers in the same format, using the same node categories and edge predicates. Because of this, any KP that speaks TRAPI can be plugged into the ARAX_expander module with ease. In fact, ARAX_expander (Sec. 2.1.1) dynamically selects TRAPI KPs that are registered in the SmartAPI registry [31], meaning ARAX is able to add new KPs without any human intervention.

##### ARAX Node Synonymizer

Because there are many overlapping controlled vocabularies within biomedicine, concepts (or the nodes that represent them) can often be described using multiple identifiers. For instance, MONDO:0019391, DOID:13636, and ORPHANET:84 are all valid identifiers for the disease Fanconi anemia, coming from the Monarch Disease Ontology [40], the Disease Ontology [41], and the Orphanet Rare Disease Ontology [42, 43], respectively. Different knowledge providers may refer to the same concept using different identifiers, making it challenging to integrate their results. To address this problem, we created the ARAX Node Synonymizer, a node synonym mapping service built into ARAX. The Node Synonymizer determines node equivalencies by combining four kinds of evidence:

1. Concept equivalence information provided by a Translator web service called the Standards and Reference Implementation Node Normalizer^9^,
2. Nodes connected by biolink:same_as relations in RTX-KG2,
3. Nodes with identical names, and
4. Node semantic type compatibility.

The Node Synonymizer uses this information to partition a given set of node identifiers into sets of semantically equivalent identifiers. Each of these clusters is assigned a single representative concept identifier from among the cluster’s members. ARAX uses the Node Synonymizer to map all node identifiers returned from knowledge providers to these canonical identifiers, facilitating merging of the knowledge providers’ responses.

### 2.3 Interfaces for accessing ARAX

#### 2.3.1 ARAX Web Browser Interface

The ARAX Web Browser User Interface (“ARAX UI”; arax.rtx.ai) provides an intuitive, human-friendly mechanism for querying ARAX and exploring the answers it returns. It allows users to formulate biomedical questions in four different formats: an interactive visual query graph builder, TRAPI Operations, TRAPI JSON, and ARAXi. The interactive query graph builder allows users to construct a query graph in a visual fashion via drop-down lists and clickable buttons, with no JSON or TRAPI knowledge required. The TRAPI Operations input aids users in creating a TRAPI-compliant Workflow by providing a drop-down list of allowable Operations and guided entry of Operation parameters. The TRAPI JSON input method facilitates sharing and re-use of graph analysis workflows. The ARAXi input method (described in Section 2.1) provides users with easy-to-understand syntax to formulate more complex query workflows, along with drop-down menus of ARAXi commands to facilitate discovery. For each query posted to the ARAX UI, the output includes five sections:

1. **Summary**: Summarizes the answers to the query in a simple table sorted by ARAX-defined scores that can be considered a measure of confidence (Sec. 2.1.5).
2. **Knowledge Graph**: Provides a visual representation of the answer KG for easy viewing of its topology.
3. **Results**: Provides a display of all the results that satisfy the input query graph, ranked as in Section 2.1.5. Clicking on individual results displays an interactive graphical representation of the result. Within each result, clicking on nodes and edges displays detailed information including evidence/provenance information and concept descriptions.
4. **Messages**: Shows all messages logged during processing of the query, including any errors or warnings.
5. **Provenance**: Shows the number of edges in the final KG by predicate type and knowledge source(s) that they came from. (May require refreshing the Web browser page to display data.)

The ARAX UI provides persistent URLs that enable fast retrieval of results from previously run queries. Apart from these main functions, the ARAX UI also provides three additional tools/services:

- **Synonym Lookup**: Exposes the ARAX Node Synonymizer (described in “ARAX Node Synonymizer” in Section 2.2.2); a user can input a concept identifier or name and see its set of equivalent identifiers and other synonym information.
- **Dev Info**: Provides a JSON-formatted view of UI-server communications for development purposes.
- **System Activity**: Displays the status and activity of previous queries submitted to ARAX over a user-selectable time interval and retrieves corresponding queries and results (if they completed successfully).

#### 2.3.2 ARAX API

ARAX can also be used via its publicly-accessible API^10^, which is listed in the SmartAPI registry [31]. The ARAX API includes /`query` and /`asyncquery` endpoints that accept TRAPI input, as well as an /entity endpoint that exposes the ARAX Node Synonymizer service, allowing users to programmatically retrieve equivalent identifiers and other synonym information for a given concept.

## 3 Use Cases

To illustrate how ARAX can be used to explore biomedical knowledge, we present three different use cases: two focused queries, one to search for molecular mechanisms in bipolar disorder and one to search for the mechanism of action for an antiviral for COVID-19 disease, and a larger-scale analysis concerning drug repurposing.

### 3.1 Use Case 1: Bipolar disorder

Bipolar disorder is a mental health condition that affects over 46M people worldwide [44]. It is characterized by mood swings that can include excessive happiness (mania) and/or excessive sadness (depression). The causes of bipolar disorder are not well understood, but are thought to include both genetic and environmental factors, with stress and substance abuse contributing to disease severity. It has been recently proposed [45] that decreased expression of D-amino acid oxidase (DAO) in cerebellar neurons might increase the risk for bipolar disorder by affecting the regulation of N-methyl-D-aspartate receptors (NMDARs). Using the ARAX web browser interface, we explored the connection between DAO and bipolar disorder via protein intermediaries in the context of a two-hop query graph (e.g., “DAO—*—bipolar disorder” where * represents a peptide/protein mediator). Within the browser interface, we posted the ARAXi commands of a two-hop query graph to ARAX, which returned 17 result subgraphs (Fig. 4). The query and its results are available as online supplementary material.^11^ The “N-methyl-D-aspartate receptors” result (which we expected to see based on prior literature [45]) is in the top 10 highest-scoring results, based on ARAX’s result-graph ranking algorithm (see Sec. 2.1.5). For some of the other peptide/protein results returned, ARAX provided hyperlinks to publications that support the relevant relations in the specific DAO-to-bipolar result graph such as the amino acid proline (Fig. 5).

**Figure 4:**
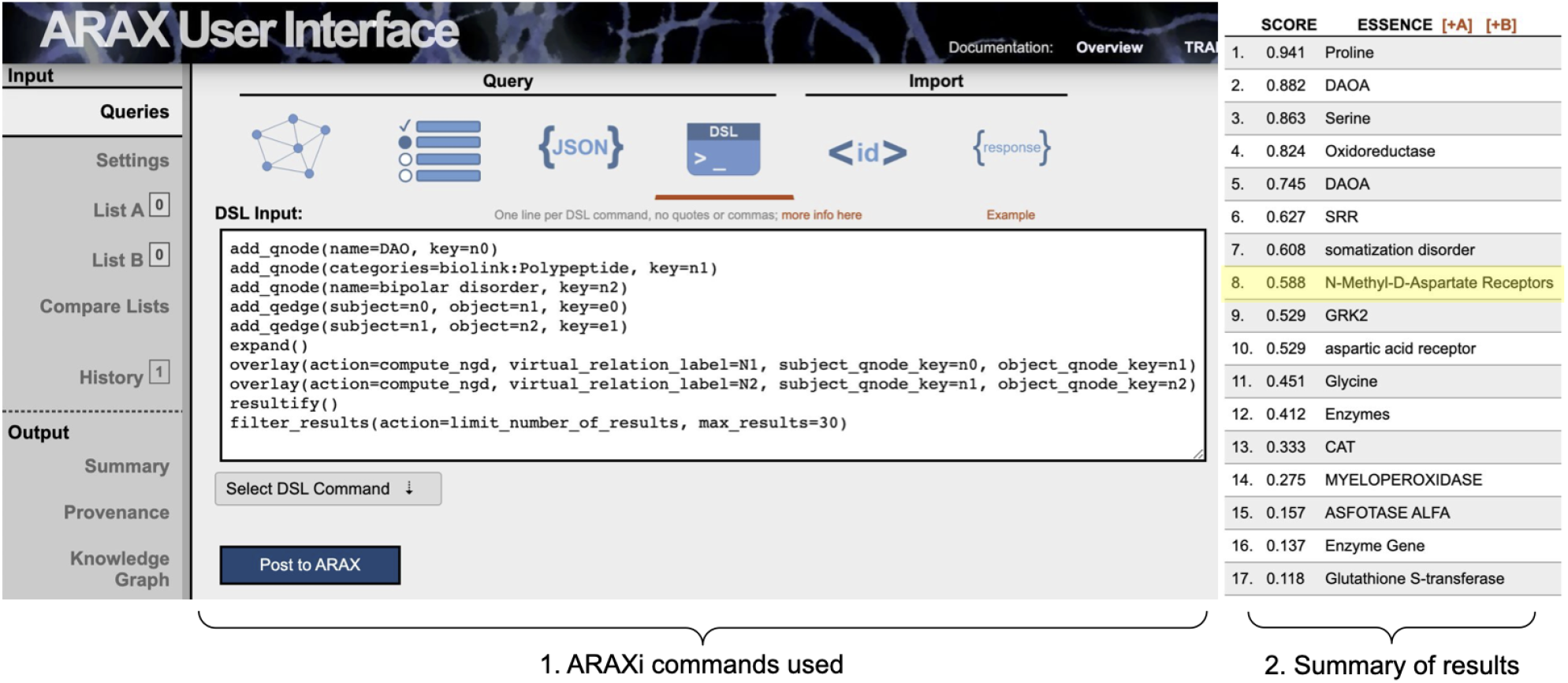
ARAX Web UI showing the ARAXi workflow (1) used for Use Case 1 (exploring what proteins might be involved in the connection between DAO and bipolar disorder) and the 17 returned results with their corresponding ranking scores (2), as they appear in the Summary tab in the UI. The expected result of “N-methyl-D-aspartate receptors” is highlighted in yellow.

**Figure 5:**
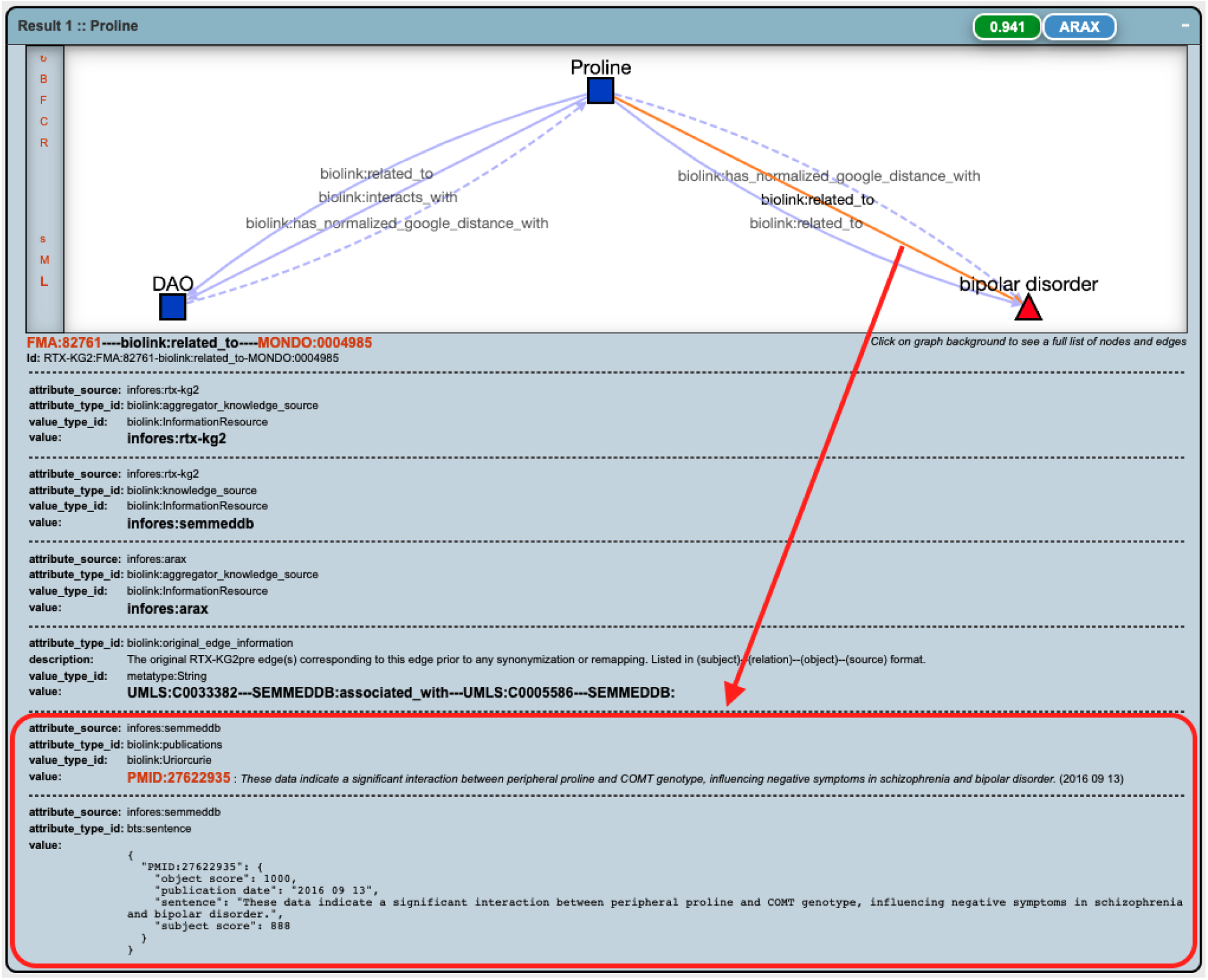
A “result graph” for a single result (“proline”) for the two-hop query for mediators between DAO and bipolar disorder, with supporting publication (PMID:27622935) and its corresponding relevant sentence shown (highlighted by a red square) for a specific edge “Proline—biolink:related_to—bipolar disorder”.

### 3.2 Use Case 2: Coronavirus disease 2019

Coronavirus disease 2019 (COVID-19), caused by the SARS-CoV-2 pathogen, has resulted in a worldwide pandemic. Researchers have made extensive efforts [46, 47, 48, 49, 50] to repurpose existing drugs to treat COVID-19. Remdesivir was the first repurposed antiviral drug that was approved by the United States Food and Drug Administration for the treatment of COVID-19. Remdesivir was originally developed to treat hepatitis C and then found to be effective against many other viral diseases including Ebola and other coronavirus diseases. Its mechanism of action is to interfere with the viral RNA-dependent RNA polymerase (RdRp) [51]. By using ARAX, we can provide a series of potential three-hop biological-relation paths between remdesivir and COVID-19 via RNA-directed RNA polymerase. Figure 6 shows the top 25 ARAX results with the corresponding ARAXi commands. The query and its results are available as online supplementary material.^12^ A top-ranking result, “Virus Replication”, is precisely the mechanism of action that has been reported by various researchers [52, 53, 51]. Furthermore, ARAX results can reveal potential downstream mediators of therapeutic efficacy such as release of type I interferon, an antiviral cytokine which SARS-CoV-2 infection is known to inhibit [54] (see result “IFNB1” for this query).

**Figure 6:**
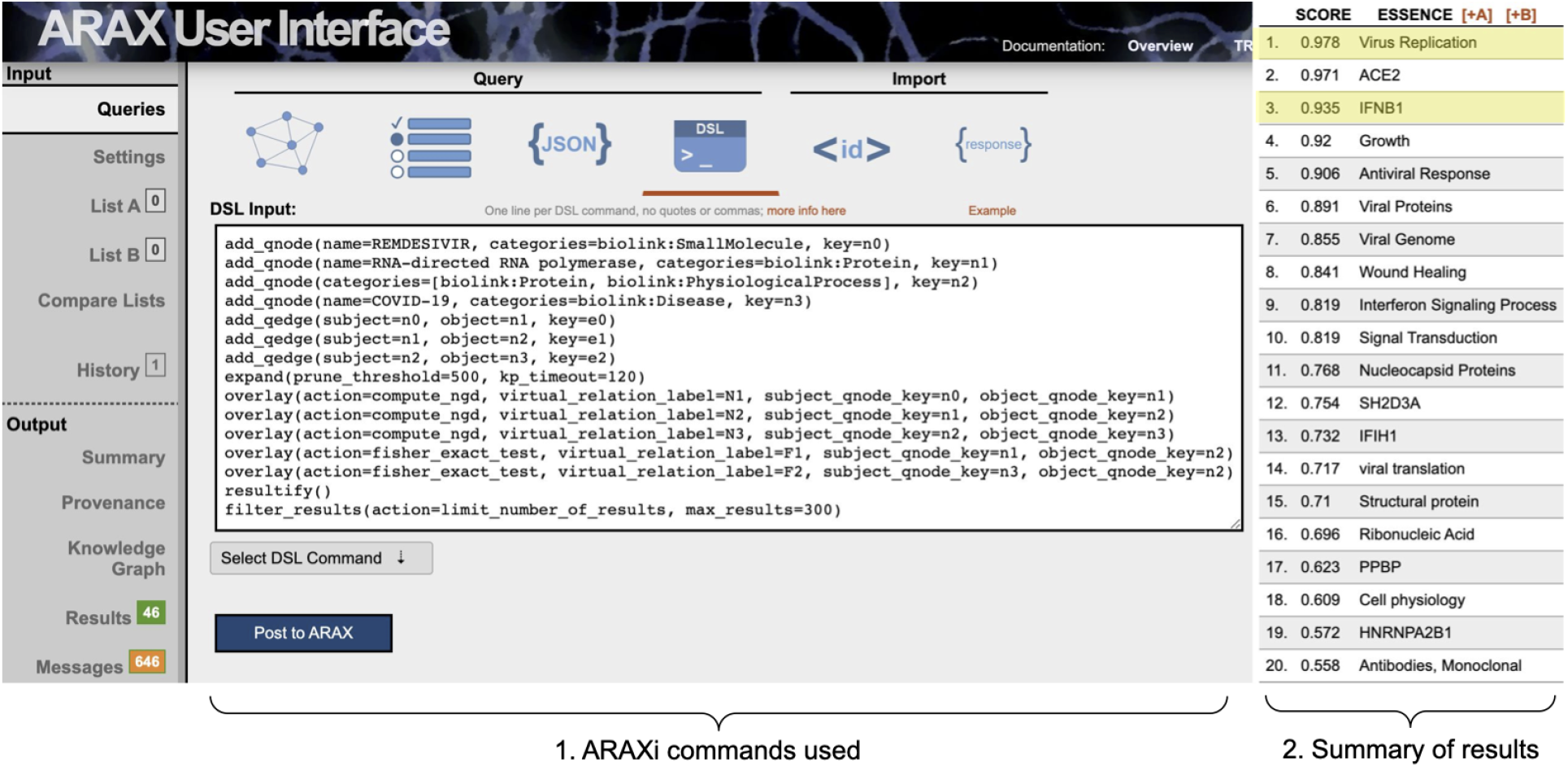
ARAXi workflow (1) for finding the remdesivir mechanism-of-action downstream of RdRp for treating COVID-19 (Use Case 2), along with the top 25 returned results (2). The results of “Virus Replication” and “IFNB1” mentioned in the main text are highlighted in yellow.

### 3.3 Use Case 3: Recovering drug/disease relationships

Drug repurposing is a strategy for finding treatments for diseases by searching for and validating new indications for existing (i.e., already approved) drugs. Computational approaches to this task often involve developing a link-prediction model: a model that, when given a drug and disease, predicts the probability that the drug treats the disease. One such model, given in Section 2.1.2, is accessible via ARAX. Here, we describe the performance of that model in predicting “treats” edges in the RTX-KG2 KP. Importantly, this model was not trained on RTX-KG2 “treats” edges but rather data from SemMedDB [6], MyChem [55], and NDF-RT [56] (as described in [33]), though some RTX-KG2 “treats” edges may exist in these three training datasets^13^. In RTX-KG2, there are 266k `biolink:treats` edges. We used the model in Section 2.1.2 to predict, for all such 266K drug/disease^14^ pairs, the probability that a “treats” relationship exists between them. We then selected all (drug, 〈`“non-treats” relation`〉, disease) triples, where `“non-treats” relation` is any relation besides the Biolink predicates: `treats, prevents, treated_by, ameliorates, is_ameliorated_by, related_to`, or `disrupts`. We used the model to calculate the link prediction probabilities of all 184k such “non-treats” edges. Figure 7 shows the distributions of this link prediction task, demonstrating the model can successfully distinguish between drug/disease pairs with the “treats” or “not-treats” relationship. Using a cutoff probability of 0.8, 40% of existing RTX-KG2 “treats” relationships were recovered, while only 17% of “non-treats” edges were above this cutoff.

**Figure 7:**
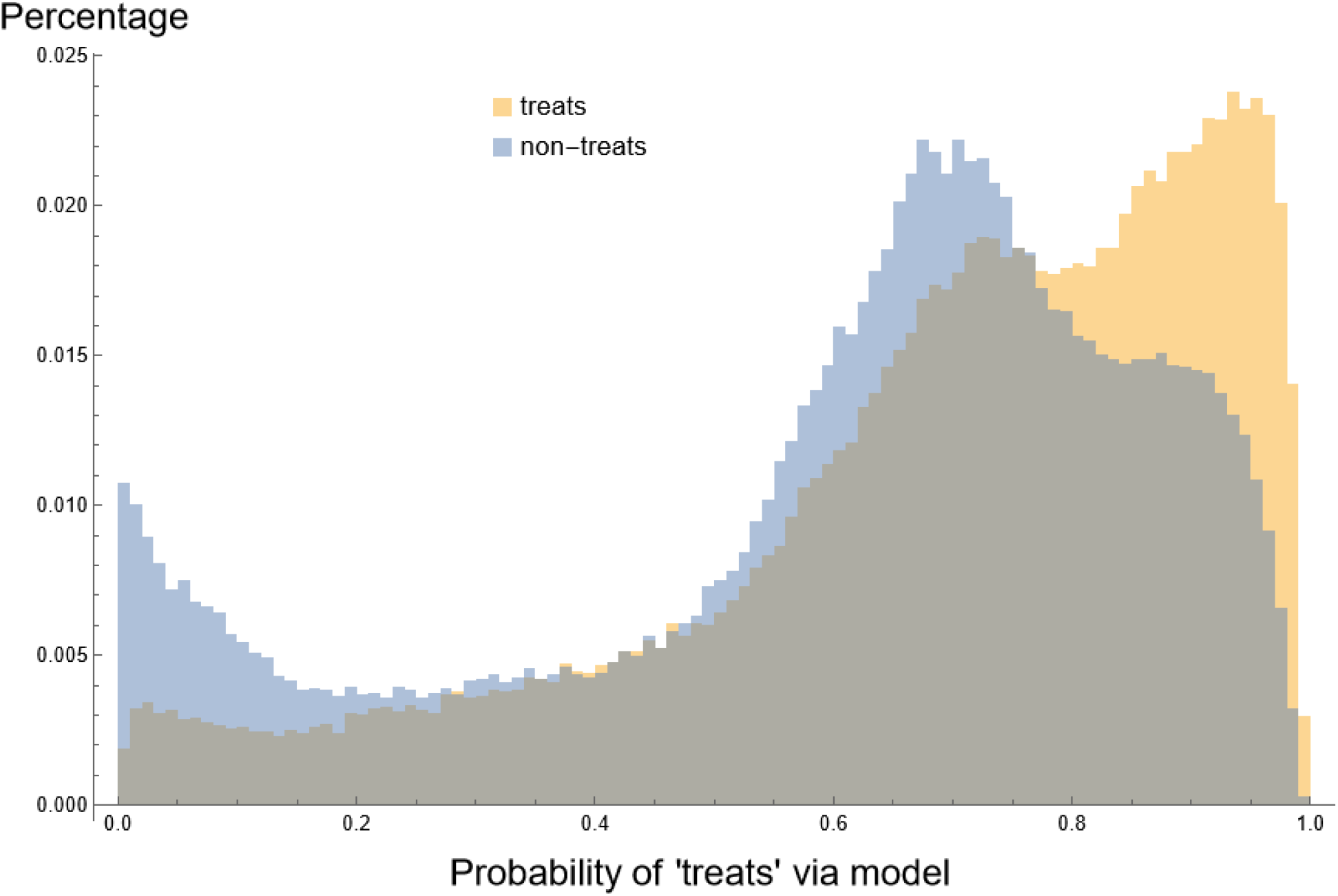
Overlaid histograms of the link prediction probability for the “treats” relationship for all drug/disease pairs having a `biolink:treats` edge between them (in yellow) or non-treats edge between them (in blue).

## 4 Discussion

The purpose of ARAX and Translator is to enable integrated analysis of structured biomedical knowledge in order to provide ranked, coherent answers to translational biomedical questions. ARAX’s three key innovations—ARAXi, its approximately 40 knowledge providers, and its result subgraph ranking algorithm—together provide significant leverage to enable a researcher to tackle translational questions that are more complex and rely more on basic science (and not just on clinically-validated) knowledge. Further, ARAX’s web browser interface enables composing queries and browsing/exploring results, without having to programmatically post-process results or query the API.

While it shares the goal of leveraging knowledge to advance translation, ARAX is not an AI-based Biomedical Question-Answering system such as IBM Watson [57], MedQA [58], or BioSQUASH [59]. Such question-answering systems can provide a direct semantic answer computationally extracted from relevant biomedical documents via natural language processing (NLP) techniques. Thus, the knowledge used in these systems is only limited to document-based knowledge. Further, ARAX differs from the NLP-based document search tool BioMed Explorer^15^; BioMed Explorer is designed to answer a question by finding relevant resources (such as articles) using semantic search and showing excerpts from those resources. ARAX, in contrast, is based on multiple types of data sources (e.g., publications, databases, electronic health records, etc.). Instead of directly providing an intuitive answer, ARAX is more like a “searching & computing engine” where it provides a self-developed domain language to translate a specific biomedical question into a query graph, “search” for answers from different knowledge providers, and then perform computations with the resulting graphs. Thus, it can facilitate querying and exploring a significant fraction of published biomedical knowledge and public biomedical knowledge-bases. However, unlike search engines or semantic search engines, ARAX allows users to create custom knowledge graph analysis workflows, as well as access reasoning modules for specific translational applications like drug repurposing.

ARAX was built to help biomedical researchers explore structured knowledge and help generate new hypotheses. ARAX is for research purposes and is not meant to be used by clinicians in the course of treating patients. As the system is being actively developed, there is no expectation that results from queries run on ARAX will be retained indefinitely. Future enhancements to ARAX will include increasing the Node Synonymizer’s accuracy and improving drug-treats-disease prediction via alternative embedding models and classifiers.

## 5 Conclusion

ARAX is a computational reasoning tool that allows users to easily extract, explore, and analyze knowledge from diverse biomedical resources. With self-developed ARAXi, ARAX allows users to encode their complex biomedical questions into specific KG analysis workflows in an intuitive manner. These workflows can intelligently integrate data from multiple biomedical knowledge providers, analyze and filter that data, and rank the final results. The ARAX UI further simplifies this process by providing multiple query options and facilitating interactive exploration of query results. The ARAX API enables incorporating ARAX’s knowledge retrieval and reasoning capabilities within a workflow involving non-ARAX tools. We believe that ARAX can help life sciences researchers to more effectively interpret new research findings and develop new hypotheses.

## Acknowledgements

We thank Mark Williams, Tyler Beck, Noel Southall, Christine Colvis, Sarah Stemann, Debbi Adelakun, Melissa Haendel, Chris Bizon, Karamarie Fecho, Patrick Wang, Chris Mungall, Sierra Moxon, Matt Brush, Paul Shannon, Andrew Su, Chunlei Wu, Kevin Xin, Will Byrd, Andrew Crouse, John Osborne, Jeff Henrikson, Greg Rosenblatt, Sergio Baranzini, Matt Might, Arnab Nandi, Sui Huang, and Liang Huang for advice and/or feedback on ARAX. We thank Dac-Trung Nguyen, Deqing Qu, Tim Yoon, Tim Putman, Richard Bruskiewicz, Pouyan Ahmadi, Ujjval Kumaria, Jason McClelland, Yao Yao, Zheng Liu, David Palzer, Sarah Oliphant, Boris Cao, Ke Wang, Max Wang, Jamie Slome, Kanna Bhargav Chevva, and Daniel Lin for technical assistance.

## Abbreviations

KG: knowledge graph
KP: knowledge provider
TRAPI: Translator Reasoner application programming interface

## Footnotes

1 neo4j.com/developer/cypher

2 github.com/NCATSTranslator/ReasonerAPI

3 github.com/biolink/biolink-model

4 github.com/NCATSTranslator/OperationsAndWorkflows

5 smart-api.info/registry

6 spoke.ucsf.edu/data-tools

7 github.com/biothings/biothings_explorer

8 The only exceptions are two in-house KPs that are stored locally and queried using SQL: Drug-Treats-Disease and Normalized PubMed Distance (Sec. 2.1.2).

9 github.com/TranslatorSRI/NodeNormalization

10 arax.rtx.ai/api/arax/v1.3/openapi.json

11 [github:RTXteam/RTX/notes/vignettes.md#bipolar-disorder](https://github.com/RTXteam/RTX/blob/master/notes/vignettes.md#bipolar-disorder)

12 github:RTXteam/RTX/notes/vignettes.md#covid-19

13 Namely, those edges from SemMedDB, which is one of RTX-KG2’s sources; MyChem and RTX-KG2 also have some overlapping sources, such as DrugCentral.

14 Note here that “drug” means `biolink:Drug or biolink:SmallMolecule`, and `“disease” means biolink:Disease` or `biolink:PhenotypicFeature`.

15 sites.research.google/biomedexplorer/

## References

[1] David Mendez, Anna Gaulton, A Patrícia Bento, Jon Chambers, Marleen De Veij, Eloy Félix, María Paula Magariños, Juan F Mosquera, Prudence Mutowo, Michal Nowotka, María Gordillo-Marañón, Fiona Hunter, Laura Junco, Grace Mugumbate, Milagros Rodriguez-Lopez, Francis Atkinson, Nicolas Bosc, Chris J Radoux, Aldo Segura-Cabrera, Anne Hersey, and Andrew R Leach. ChEMBL: towards direct deposition of bioassay data. Nucleic Acids Research, 47(D1):D930–D940, January 2019.

[2] David S Wishart, Craig Knox, An Chi Guo, Savita Shrivastava, Murtaza Hassanali, Paul Stothard, Zhan Chang, and Jennifer Woolsey. DrugBank: a comprehensive resource for in silico drug discovery and exploration. Nucleic Acids Research, 34(Database issue):D668–72, January 2006.

[3] Antonio Fabregat, Florian Korninger, Guilherme Viteri, Konstantinos Sidiropoulos, Pablo Marin-Garcia, Peipei Ping, Guanming Wu, Lincoln Stein, Peter D’Eustachio, and Henning Hermjakob. Reactome graph database: Efficient access to complex pathway data. PLoS Comput. Biol., 14(1):e1005968, January 2018.

[4] Barry Smith, Michael Ashburner, Cornelius Rosse, Jonathan Bard, William Bug, Werner Ceusters, Louis J Goldberg, Karen Eilbeck, Amelia Ireland, Christopher J Mungall, Neocles Leontis, Philippe Rocca-Serra, Alan Ruttenberg, Susanna-Assunta Sansone, Richard H Scheuermann, Nigam Shah, Patricia L Whetzel, Suzanna Lewis, and The OBI Consortium. The OBO Foundry: coordinated evolution of ontologies to support biomedical data integration. Nature Biotechnology, 25(11):1251–1255, 2007.

[5] M Kanehisa and S Goto. KEGG: Kyoto encyclopedia of genes and genomes. Nucleic Acids Research, 28(1): 27–30, January 2000.

[6] Halil Kilicoglu, Dongwook Shin, Marcelo Fiszman, Graciela Rosemblat, and Thomas C. Rindflesch. Semmeddb: a pubmed-scale repository of biomedical semantic predications. Bioinformatics, 28(23):3158–3160, 2012. ISSN 1367-4803. doi:10.1093/bioinformatics/bts591.

[7] UniProt Consortium. UniProt: the universal protein knowledgebase in 2021. Nucleic Acids Research, 49(D1): D480–D489, January 2021.

[8] Olivier Bodenreider. The Unified Medical Language System (UMLS): integrating biomedical terminology. Nucleic Acids Research, 32(Database issue):D267–70, January 2004.

[9] John F Sowa. Conceptual graphs as a universal knowledge representation. Comput. Math. Appl., 23(2-5):75–93, January 1992.

[10] M Joubert, M Fieschi, and F Volot. Review of biomedical knowledge and data representation with conceptual graphs. Methods Inf. Med., 37(01):86–96, 1998.

[11] M Joubert, M Fieschi, J J Robert, F Volot, and D Fieschi. UMLS-based conceptual queries to biomedical information databases: an overview of the project ARIANE. unified medical language system. J. Am. Med. Inform. Assoc., 5(1):52–61, January 1998.

[12] Michel Dumontier, Alison Callahan, Jose Cruz-Toledo, Peter Ansell, Vincent Emonet, François Belleau, and Arnaud Droit. Bio2RDF release 3: a larger connected network of linked data for the life sciences. In Proceedings of the 2014 International Conference on Posters & Demonstrations Track, volume 1272, pages 401–404. Citeseer, 2014.

[13] Janet Piñero, Àlex Bravo, Núria Queralt-Rosinach, Alba Gutiérrez-Sacristán, Jordi Deu-Pons, Emilio Centeno, Javier García-García, Ferran Sanz, and Laura I. Furlong. DisGeNET: a comprehensive platform integrating information on human disease-associated genes and variants. Nucleic Acids Research, 45(D1):D833–D839, October 2016. doi:10.1093/nar/gkw943. URL https://doi.org/10.1093/nar/gkw943.

[14] Daniel Scott Himmelstein, Antoine Lizee, Christine Hessler, Leo Brueggeman, Sabrina L Chen, Dexter Hadley, Ari Green, Pouya Khankhanian, and Sergio E Baranzini. Systematic integration of biomedical knowledge prioritizes drugs for repurposing. eLife, 6, September 2017. doi:10.7554/elife.26726. URL https://doi.org/10.7554/elife.26726.

[15] Antonio Messina, Haikal Pribadi, Jo Stichbury, Michelangelo Bucci, Szymon Klarman, and Alfonso Urso. BioGrakn: A knowledge graph-based semantic database for biomedical sciences. In Advances in Intelligent Systems and Computing, Advances in intelligent systems and computing, pages 299–309. Springer International Publishing, Cham, 2018.

[16] Kenneth Morton, Patrick Wang, Chris Bizon, Steven Cox, James Balhoff, Yaphet Kebede, Karamarie Fecho, and Alexander Tropsha. ROBOKOP: an abstraction layer and user interface for knowledge graphs to support question answering. Bioinformatics, 35(24):5382–5384, August 2019. doi:10.1093/bioinformatics/btz604. URL https://doi.org/10.1093/bioinformatics/btz604.

[17] Geoffrey Sanders, Roger Pearce, and Sergio E. Baranzini. Topological analysis of the SPOKE graph. Technical report, U. S. Department of Energy, 9 2020. URL https://www.osti.gov/biblio/1669224. DOI:10.2172/1669224.

[18] Vassilis N. Ioannidis, Xiang Song, Saurav Manchanda, Mufei Li, Xiaoqin Pan, Da Zheng, Xia Ning, Xiangxiang Zeng, and George Karypis. DRKG - drug repurposing knowledge graph for COVID-19. Web page:https://github.com/gnn4dr/DRKG/, 2020.

[19] E C Wood, Amy K Glen, Lindsey G Kvarfordt, Finn Womack, Liliana Acevedo, Timothy S Yoon, Chunyu Ma, Veronica Flores, Meghamala Sinha, Yodsawalai Chodpathumwan, Arash Termehchy, Jared C Roach, Luis Mendoza, Andrew S Hoffman, Eric W Deutsch, David Koslicki, and Stephen A Ramsey. RTX-KG2: a system for building a semantically standardized knowledge graph for translational biomedicine. bioRxiv preprint biorxiv:464747, October 2021. doi:10.1101/2021.10.17.464747. URL https://doi.org/10.1101/2021.10.17.464747.

[20] Deepak R Unni, Sierra AT Moxon, Michael Bada, Matthew Brush, Richard Bruskiewich, J Harry Caufield, Paul A Clemons, Vlado Dancik, Michel Dumontier, Karamarie Fecho, et al. Biolink Model: A universal schema for knowledge graphs in clinical, biomedical, and translational science. Clinical and Translational Science, 2022.

[21] Scott Wen-tau Yih, Ming-Wei Chang, Xiaodong He, and Jianfeng Gao. Semantic parsing via staged query graph generation: Question answering with knowledge base. In Proceedings of the Joint Conference of the 53rd Annual Meeting of the ACL and the 7th International Joint Conference on Natural Language Processing of the AFNLP, 2015.

[22] Renzo Angles and Claudio Gutierrez. The expressive power of sparql. In International Semantic Web Conference, pages 114–129. Springer, 2008.

[23] Aaron Birkland and Golan Yona. BIOZON: a system for unification, management and analysis of heterogeneous biological data. BMC Bioinformatics, 7(1), February 2006. doi:10.1186/1471-2105-7-70. URL https://doi.org/10.1186/1471-2105-7-70.

[24] Antonio Messina, Antonino Fiannaca, Laura La Paglia, Massimo La Rosa, and Alfonso Urso. BioGraph: a web application and a graph database for querying and analyzing bioinformatics resources. BMC SystemsBiology, 12(S5), November 2018. doi:10.1186/s12918-018-0616-4. URL https://doi.org/10.1186/s12918-018-0616-4.

[25] Charlotte A. Nelson, Atul J. Butte, and Sergio E. Baranzini. Integrating biomedical research and electronic health records to create knowledge-based biologically meaningful machine-readable embeddings. Nature Communications, 10(1), July 2019. doi:10.1038/s41467-019-11069-0. URL https://doi.org/10.1038/s41467-019-11069-0.

[26] Jiwen Xin, Cyrus Afrasiabi, Sebastien Lelong, Andrew I. Su, and Chunlei Wu. Biothings Explorer: utilizing JSON-LD for linking biological APIs to facilitate knowledge discovery, 2017. URL https://f1000research.com/posters/6-1173.

[27] William E. Byrd, Gregory Rosenblatt, Michael John Patton, Thi K. Tran-Nguyen, Marissa Zheng, Apoorv Jain, Michael Ballantyne, Katherine Zhang, Mei-Jan Chen, Jordan Whitlock, Mary E. Crumbley,, Jillian Tinglin, Kaiwen He, Yizhou Zhang, Jeremy D. Zucker, Joseph A. Cottam, Nada Amin, John Osborne, Andrew Crouse, and Matthew Might. mediKanren: A system for bio-medical reasoning. In Proceedings of the 2020 ACM SIGPLAN International Conference on Functional Programming, 2020.

[28] Christopher P. Austin. Translating translation. Nature Reviews Drug Discovery, 17(7):455–456, April 2018. doi:10.1038/nrd.2018.27. URL https://doi.org/10.1038/nrd.2018.27.

[29] Translator Consortium. Toward a universal biomedical data translator. Clinical and Translational Science, 12(2): 86–90, 2019.

[30] Translator Consortium. The Biomedical Data Translator program: Conception, culture, and community. Clinical and Translational Science, 12(2):91–94, March 2019. doi:10.1111/cts.12592. URL https://doi.org/10.1111/cts.12592.

[31] Amrapali Zaveri, Shima Dastgheib, Chunlei Wu, Trish Whetzel, Ruben Verborgh, Paul Avillach, Gabor Korodi, Raymond Terryn, Kathleen Jagodnik, Pedro Assis, and Michel Dumontier. smartAPI: Towards a more intelligent network of Web APIs. In Eva Blomqvist, Diana Maynard, Aldo Gangemi, Rinke Hoekstra, Pascal Hitzler, and Olaf Hartig, editors, The Semantic Web, pages 154–169, Cham, 2017. Springer International Publishing. ISBN 978-3-319-58451-5.

[32] Pauli Virtanen, Ralf Gommers, Travis E. Oliphant, Matt Haberland, Tyler Reddy, David Cournapeau, Evgeni Burovski, Pearu Peterson, Warren Weckesser, Jonathan Bright, Stéfan J. van der Walt, Matthew Brett, Joshua Wilson, K. Jarrod Millman, Nikolay Mayorov, Andrew R. J. Nelson, Eric Jones, Robert Kern, Eric Larson, C J Carey, İlhan Polat, Yu Feng, Eric W. Moore, Jake VanderPlas, Denis Laxalde, Josef Perktold, Robert Cimrman, Ian Henriksen, E. A. Quintero, Charles R. Harris, Anne M. Archibald, Antônio H. Ribeiro, Fabian Pedregosa, Paul van Mulbregt, and SciPy 1.0 Contributors. SciPy 1.0: Fundamental Algorithms for Scientific Computing in Python. Nature Methods, 17:261–272, 2020. doi:10.1038/s41592-019-0686-2.

[33] Finn Womack, Jason McClelland, and David Koslicki. Leveraging distributed biomedical knowledge sources to discover novel uses for known drugs. bioRxiv preprint biorxiv:765305, September 2019. doi:10.1101/765305. URL https://doi.org/10.1101/765305.

[34] Aditya Grover and Jure Leskovec. node2vec: Scalable feature learning for networks, 2016. URL https://arxiv.org/abs/1607.00653.

[35] William L. Hamilton, Rex Ying, and Jure Leskovec. Inductive representation learning on large graphs, 2017. URL https://arxiv.org/abs/1706.02216.

[36] Rudi Cilibrasi and Paul M. B. Vitanyi. The Google similarity distance. arXiv preprint arxiv:cs/0412098, 2004. doi:10.48550/ARXIV.CS/0412098. URL https://arxiv.org/abs/cs/0412098.

[37] FB Rogers. Medical subject headings. Bulletin of the Medical Library Association, 51(1):114–116, January 1963.

[38] Casey N. Ta, Michel Dumontier, George Hripcsak, Nicholas P. Tatonetti, and Chunhua Weng. Columbia Open Health Data, clinical concept prevalence and co-occurrence from electronic Data, 5(1), November 2018. doi:10.1038/sdata.2018.273. URL https://doi.org/10.1038/sdata.2018.273.

[39] Karamarie Fecho, Emily Pfaff, Hao Xu, James Champion, Steve Cox, Lisa Stillwell, David B Peden, Chris Bizon, Ashok Krishnamurthy, Alexander Tropsha, and Stanley C Ahalt. A novel approach for exposing and sharing clinical data: the translator integrated clinical and environmental exposures service. Journal of the American Medical Informatics Association, 26(10):1064–1073, April 2019. doi:10.1093/jamia/ocz042. URL https://doi.org/10.1093/jamia/ocz042.

[40] Christopher J Mungall, Julie A McMurry, Sebastian Köhler, James P Balhoff, Charles Borromeo, Matthew Brush, Seth Carbon, Tom Conlin, Nathan Dunn, Mark Engelstad, et al. The Monarch initiative: an integrative data and analytic platform connecting phenotypes to genotypes across species. Nucleic Acids Research, 45(D1): D712–D722, 2017.

[41] Lynn M Schriml, Elvira Mitraka, James Munro, Becky Tauber, Mike Schor, Lance Nickle, Victor Felix, Linda Jeng, Cynthia Bearer, Richard Lichenstein, et al. Human disease ontology 2018 update: classification, content and workflow expansion. Nucleic Acids Research, 47(D1):D955–D962, 2019.

[42] S S Weinreich, R Mangon, J J Sikkens, M E en Teeuw, and M C Cornel. Orphanet: a European database for rare diseases. Nederlands Tijdschrift voor Geneeskunde, 152(9):518–9, 2008. ISSN 0028-2162.

[43] Drashtti Vasant, Laetitia Chanas, James Malone, Marc Hanauer, Annie Olry, Simon Jupp, Peter N Robinson, Helen Parkinson, and Ana Rath. Ordo: an ontology connecting rare disease, epidemiology and genetic data. In Proceedings of ISMB, volume 30, 2014.

[44] Alize J Ferrari, Emily Stockings, Jon-Paul Khoo, Holly E Erskine, Louisa Degenhardt, Theo Vos, and Harvey A Whiteford. The prevalence and burden of bipolar disorder: findings from the Global Burden of Disease Study 2013. Bipolar disorders, 18(5):440–450, 2016.

[45] Naushaba Hasin, Lace M. Riggs, Tatyana Shekhtman, Justin Ashworth, Robert Lease, Rediet T. Oshone, Elizabeth M. Humphries, Judith A. Badner, Pippa A. Thompson, David C. Glahn, David W. Craig, Howard J. Edenberg, Elliot S. Gershon, Francis J. McMahon, John I. Nurnberger, Peter P. Zandi, John R. Kelsoe, Jared C. Roach, Todd D. Gould, and Seth A. Ament. A rare variant in d-amino acid oxidase implicates NMDA receptor signaling and cerebellar gene networks in risk for bipolar disorder. medRxiv preprint medrxiv:2021.06.02.21258261, June 2021. doi:10.1101/2021.06.02.21258261. URL https://doi.org/10.1101/2021.06.02.21258261.

[46] Kelly Ansems, Felicitas Grundeis, Karolina Dahms, Agata Mikolajewska, Volker Thieme, Vanessa Piechotta, Maria-Inti Metzendorf, Miriam Stegemann, Carina Benstoem, and Falk Fichtner. Remdesivir for the treatment of COVID-19. Cochrane Database of Systematic Reviews, 2021(8), August 2021. doi:10.1002/14651858.cd014962. URL https://doi.org/10.1002/14651858.cd014962.

[47] Courtney Temple, Ruby Hoang, and Robert G. Hendrickson. Toxic effects from ivermectin use associated with prevention and treatment of COVID-19. New England Journal of Medicine, 385(23):2197–2198, December 2021. doi:10.1056/nejmc2114907. URL https://doi.org/10.1056/nejmc2114907.

[48] Ivan O. Rosas, Norbert Bräu, Michael Waters, Ronaldo C. Go, Bradley D. Hunter, Sanjay Bhagani, Daniel Skiest, Mariam S. Aziz, Nichola Cooper, Ivor S. Douglas, Sinisa Savic, Taryn Youngstein, Lorenzo Del Sorbo, Antonio Cubillo Gracian, David J. De La Zerda, Andrew Ustianowski, Min Bao, Sophie Dimonaco, Emily Graham, Balpreet Matharu, Helen Spotswood, Larry Tsai, and Atul Malhotra. Tocilizumab in hospitalized patients with severe COVID-19 pneumonia. New England Journal of Medicine, 384(16):1503–1516, April 2021. doi:10.1056/nejmoa2028700. URL https://doi.org/10.1056/nejmoa2028700.

[49] Soheil Hassanipour, Morteza Arab-Zozani, Bahman Amani, Forough Heidarzad, Mohammad Fathalipour, and Rudolph Martinez de Hoyo. The efficacy and safety of favipiravir in treatment of COVID-19: a systematic review and meta-analysis of clinical trials. Scientific Reports, 11(1), May 2021. doi:10.1038/s41598-021-90551-6. URL https://doi.org/10.1038/s41598-021-90551-6.

[50] Gilmar Reis, Eduardo Augusto dos Santos Moreira-Silva, Daniela Carla Medeiros Silva, Lehana Thabane, Aline Cruz Milagres, Thiago Santiago Ferreira, Castilho Vitor Quirino dos Santos, Vitoria Helena de Souza Campos, Ana Maria Ribeiro Nogueira, Ana Paula Figueiredo Guimaraes de Almeida, Eduardo Diniz Callegari, Adhemar Dias de Figueiredo Neto, Leonardo Cançado Monteiro Savassi, Maria Izabel Campos Simplicio, Luciene Barra Ribeiro, Rosemary Oliveira, Ofir Harari, Jamie I Forrest, Hinda Ruton, Sheila Sprague, Paula McKay, Alla V Glushchenko, Craig R Rayner, Eric J Lenze, Angela M Reiersen, Gordon H Guyatt, and Edward J Mills. Effect of early treatment with fluvoxamine on risk of emergency care and hospitalisation among patients with COVID-19: the TOGETHER randomised, platform clinical trial. The Lancet Global Health, 10(1):e42–e51, January 2022. doi:10.1016/s2214-109x(21)00448-4. URL https://doi.org/10.1016/s2214-109x(21)00448-4.

[51] Richard T. Eastman, Jacob S. Roth, Kyle R. Brimacombe, Anton Simeonov, Min Shen, Samarjit Patnaik, and Matthew D. Hall. Remdesivir: A review of its discovery and development leading to emergency use authorization for treatment of COVID-19. ACS Central Science, 6(5):672–683, May 2020. doi:10.1021/acscentsci.0c00489. URL https://doi.org/10.1021/acscentsci.0c00489.

[52] Goran Kokic, Hauke S. Hillen, Dimitry Tegunov, Christian Dienemann, Florian Seitz, Jana Schmitzova, Lucas Farnung, Aaron Siewert, Claudia Höbartner, and Patrick Cramer. Mechanism of SARS-CoV-2 polymerase stalling by remdesivir. Nature Communications, 12(1), January 2021. doi:10.1038/s41467-020-20542-0. URL https://doi.org/10.1038/s41467-020-20542-0.

[53] Jakob J. Malin, Isabelle Suárez, Vanessa Priesner, Gerd Fätkenheuer, and Jan Rybniker. Remdesivir against COVID- 19 and other viral diseases. Clinical Microbiology Reviews, 34(1), December 2020. doi:10.1128/cmr.00162-20. URL https://doi.org/10.1128/cmr.00162-20.

[54] Wenjing Wang, Zhuo Zhou, Xia Xiao, Zhongqin Tian, Xiaojing Dong, Conghui Wang, Li Li, Lili Ren, Xiaobo Lei, Zichun Xiang, and Jianwei Wang. SARS-CoV-2 nsp12 attenuates type I interferon production by inhibiting IRF3 nuclear translocation. Cellular & Molecular Immunology, 18(4):945–953, 2021.

[55] Jiwen Xin, Cyrus Afrasiabi, Sebastien Lelong, Julee Adesara, Ginger Tsueng, Andrew I. Su, and Chunlei Wu. Cross-linking biothings apis through json-ld to facilitate knowledge exploration. BMC Bioinformatics, 19(1):30, 2018. doi:10.1186/s12859-018-2041-5.

[56] Steven H Brown, Peter L Elkin, S Trent Rosenbloom, Casey Husser, Brent A Bauer, Michael J Lincoln, John Carter, Mark Erlbaum, and Mark S Tuttle. Va national drug file reference terminology: a cross-institutional content coverage study. Studies in health technology and informatics, 107(Pt 1):477–81, 2004. ISSN 0926-9630.

[57] D. A. Ferrucci. Introduction to “this is Watson”. IBM Journal of Research and Development, 56(3.4):1:1–1:15, May 2012. doi:10.1147/jrd.2012.2184356. URL https://doi.org/10.1147/jrd.2012.2184356.

[58] Hong Yu, Minsuk Lee, David Kaufman, John Ely, Jerome A. Osheroff, George Hripcsak, and James Cimino. Development, implementation, and a cognitive evaluation of a definitional question answering system for physicians. Journal of Biomedical Informatics, 40(3):236–251, June 2007. doi:10.1016/j.jbi.2007.03.002. URL https://doi.org/10.1016/j.jbi.2007.03.002.

[59] Zhongmin Shi, Gabor Melli, Yang Wang, Yudong Liu, Baohua Gu, Mehdi M. Kashani, Anoop Sarkar, and Fred Popowich. Question answering summarization of multiple biomedical documents. In Canadian Conference on AI, 2007.

